# Phage-encoded cationic antimicrobial peptide used for outer membrane disruption in lysis

**DOI:** 10.1101/515445

**Authors:** Ashley Holt, Jesse Cahill, Jolene Ramsey, Chandler O’Leary, Russell Moreland, Cody Martin, Don Thushara Galbadage, Riti Sharan, Preeti Sule, Kelsey Bettridge, Jie Xiao, Jeffrey Cirillo, Ryland Young

**Author notes:** co-first authors (Ashley Holt, Jesse Cahill).

## Abstract

Spanins are required for the last step in bacteriophage lysis: the disruption of the outer membrane. Bioinformatic analysis has shown that ~15% of phages lack a spanin gene, which suggests an alternate mechanism of outer membrane disruption. To address this, we selected virulent podophage ϕKT as a spaninless exemplar and tested ϕKT genes for outer membrane disruption during lysis. Hypothetical novel gene *28* causes outer membrane disruption when co-expressed with ϕKT lysis genes and complements the lysis defect of a λ spanin mutant. Gp*28* is a 56 aa cationic peptide with predicted amphipathic helical structure and is associated with the particulate fraction after lysis. Urea and KCl washes did not release gp*28* from the particulate, suggesting a strong hydrophobic interaction with the membrane. Super high-resolution microscopy supports a primarily outer membrane localization for the peptide. Additionally, holin function is not required for gp*28*-mediated lysis. Gp*28* is similar in size, charge, predicted fold, and membrane association to the human cathelicidin antimicrobial peptide LL-37. In standard assays to measure bactericidal and inhibitory effects of antimicrobial peptides on bacterial cells, synthesized gp*28* performed equivalently to LL-37. The studies presented here suggest that ϕKT Gp*28* disrupts bacterial outer membranes during lysis in a manner akin to antimicrobial peptides.

**Significance:** Here we provide evidence that ϕKT produces an antimicrobial peptide for outer membrane disruption during lysis. The disruptin is a new paradigm for phage lysis, and has no similarities to other known lysis genes. Many mechanisms have been proposed for the function of antimicrobial peptides, however there is not a consensus on the molecular basis of membrane disruption. Additionally, there is no established genetic selection system to support such studies. Therefore, the ϕKT disruptin may represent the first genetically tractable antimicrobial peptide.

## Introduction

In general, bacteriophages (phages) must lyse the host cell to release progeny (1). For phages of Gram-negative hosts, lysis requires three classes of phage proteins, each targeting one layer of the cell envelope: holins for the inner membrane (IM); endolysins for the cell wall/peptidoglycan (PG); and spanins for the outer membrane (OM) (2). In phage λ these proteins are encoded by four genes: *S* encodes the holin, *R* encodes the endolysin, and the nested genes *Rz/Rz1* encode the two spanin subunits (2, 3) (Fig. 1A). The initial steps of λ lysis have been well characterized. Lysis is initiated by the sudden formation of micron-scale holes in the IM by the holin (2, 4). These massive nonspecific holes allow the endolysin to escape to the periplasm and attack the PG (3).

**Figure 1.**
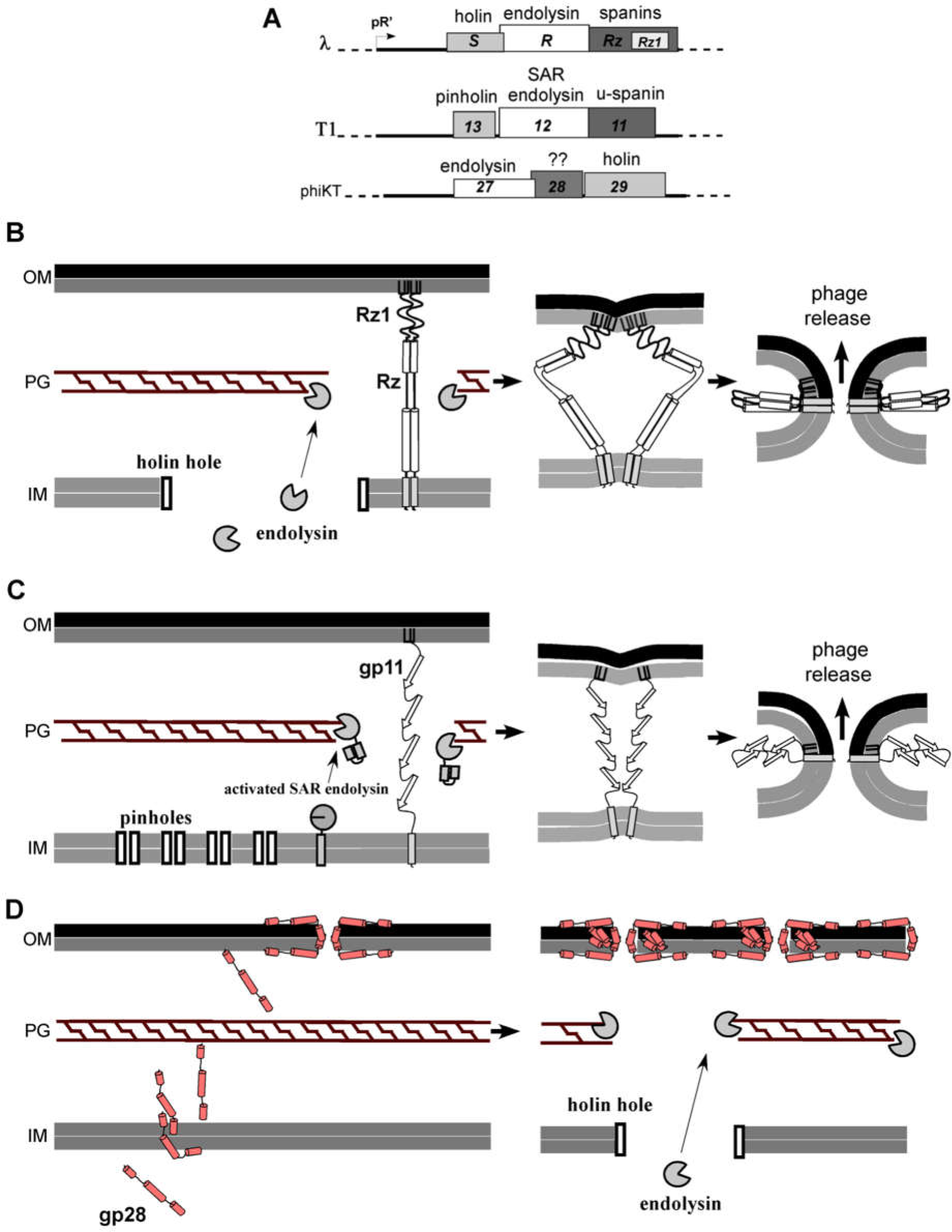
Phage use spanins or antimicrobial peptide-like proteins to disrupt the OM during lysis. A) Scaled comparisons of phages λ, T1, and ϕKT lysis cassettes. λ lysis genes are expressed under pR’ promoter control and include a classic large-hole holin, endolysin, and an embedded two-component spanin. Phage T1 encodes a pinholin-SAR endolysin system with a unimolecular spanin (u-spanin). The lysis cassette of ΦKT includes three genes: the putative endolysin (gene *27*), a hypothetical novel gene (*28*), and the putative holin (*29*). Gene *28* encodes a 56 amino acid product. B) Two-component spanin (RzRz1) lysis as observed in phage λ proceeds as the holin breaches the IM, cytoplasmic endolysin reaches and degrades the PG, and spanins are freed to associate and fuse the IM and OM, effecting phage release. C) In phage T1 lysis, the IM is compromised by a pinholin, which results in SAR endolysin release into the periplasm where it refolds and begins to degrade the PG. Finally unimolecular spanins (gp11) allow phage escape when they fuse the IM and OM. D) The disruptin model for lysis in phage ϕKT. gp*28* molecules (red) are produced with the other lysis proteins and immediately positioned in both membranes, prior to holin triggering. As gp*28* molecules enter the membrane, there is a gradual loss of membrane integrity. Holin triggering causes a hole in the IM, releasing endolysin into the periplasm. Endolysin degrades the PG, so that both the IM and PG are compromised. The OM is disrupted by gp*28*, and the cell lyses.

This holin-controlled, temporally scheduled enzymatic degradation of the PG was traditionally considered to be necessary and sufficient for lysis(1). However, recently we showed that in λ infections with spanin mutants lysis was abrogated, with the progeny virions entrapped within fragile spherical cells bounded by the intact OM(5). This dramatic phenotype had been masked in liquid culture experiments because the shearing forces attendant to aeration were sufficient to destroy these fragile cell forms and release the progeny virions.

A model for the disruption of the OM by the spanins has recently been proposed and is based on similarities with viral fusion proteins (5). The spanin genes *Rz* and *Rz1* encode subunits of a complex that connects the IM and OM (Fig. 1B)(6). Rz is a 153 aa integral membrane protein (i-spanin), whereas Rz1 is a 40 aa outer membrane lipoprotein (o-spanin)(7). Both *Rz* and *Rz1* are covalent homodimers and the complex is a heterotetramer formed by C-terminal interactions of the periplasmic domains(4, 8). The model proposes that when the PG is destroyed by the endolysin, the spanin complexes are liberated from the encaging PG meshwork and able to undergo tertiary and quaternary conformational changes leading to fusion of the IM and OM(9).

Bioinformatic analysis of 677 genomes of phages that infect Gram-negative hosts found that 586 encode identifiable spanins: 528 have two-component spanins systems encoding separate i-spanin and o-spanin subunits like λ; 58 are of a different type, the unimolecular spanins, in which a single polypeptide has both OM lipoprotein and IM transmembrane domain determinants (7, 10). The u-spanins are proposed as analogs of Class II viral fusion proteins (Fig. 1C). Interestingly, we were unable to identify spanins of either type in 91 genomes (“spaninless” phages). Since OM disruption is required for lysis(5), we wondered if these phages used a different pathway. To address this issue, we chose to examine the coliphage ΦKT from the spaninless group. The results unambiguously establish that some phages use an entirely different mode of OM disruption. The properties and function of the ΦKT protein that effects this disruption, designated as a disruptin, are discussed in terms of a new model for the coordination of phage lysis.

## Results

### ΦKT gp*28* complements a spanin defect in λ lysis

ΦKT is a virulent podophage of *E. coli* 4s, isolated from horse feces(11). Like other members of the T7 superfamily, the late genes, including cistrons required for lysis, are arranged in tandem at the right end of the genome, from ~16 kb on. It should be noted that for the endolysin gene, an incorrect downstream start codon had been chosen in the original Genbank submission (12). Once this correction was made, gp_27_ was identified as a 150 aa polypeptide with the residues Glu14-Cys23-Thr29 serving as the canonical catalytic triad motif E-X_8_-C/D-X_5_-T of the T4 endolysin family of glycoside hydrolases (13). Based on its proximity to the endolysin gene and its multiple predicted transmembrane domain architecture, gene *29* was annotated as the holin: 127 aa, 13.2 kDa, with 4 predicted transmembrane domains. A rigorous analysis showed that there were no possible lipobox sequences in any of the remaining open reading frames (10). This rules out the presence of a putative spanin of either the unimolecular or two-component type, since both involve a lipoprotein. Therefore, we wondered whether ϕKT infections ended in explosive lysis, which in all well-characterized phages of Gram-negative hosts requires a spanin for disruption of the OM. Examination of ϕKT infections of *E. coli* 4s, revealed that the infection cycles terminated in a localized and catastrophic disruption of the cell requiring on the order of one second (Fig. 2A), indistinguishable from lytic events documented for coliphage lambda(5). Moreover efficient lysis was observed irrespective of the presence of millimolar quantities of Mg^++^ cations, which stabilize the fragile spherical cells that accumulate in spanin mutant infections. We conclude that in ϕKT infections the outer membrane is disrupted by phage-encoded lysis proteins.

**Figure 2.**
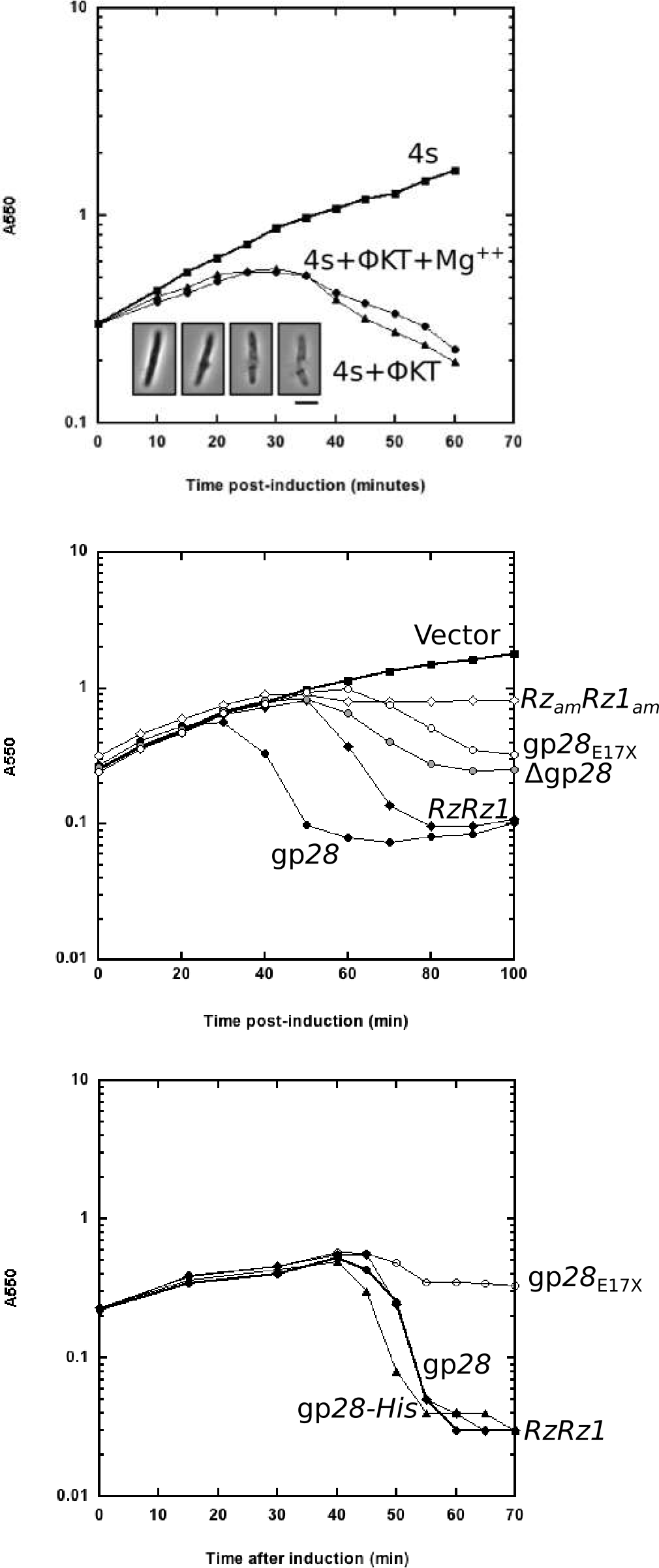
ϕKT gp*28* is necessary and sufficient for outer membrane disruption. A) At an A550 of 0.3, 4s cultures are infected with ΦKT at an MOI of ~30. After 90 min adsorption on ice (t=0), cultures are returned to 37°C, and the absorbance is monitored and plotted as a function of time for the following cultures: 4s, 4s infected by ΦKT in the presence **of** absence of magnesium ions. Inset: A single cell was imaged at the time of lysis. The montage shown spans approximately one second. Scale bar represents 5 um. B) The A550 of MG1655 cells, containing both pQ and the respective pRE plasmids, is monitored for 100 minutes. Each culture was induced with 1 mM IPTG at time=0. The pRE plasmid inserts are the ϕKT lysis cassette construct with gp*27*-gp*28*-gp*29* and the same cassette with an in-frame deletion (Δgp***28***) and truncated mutant (gp*28*_*E17X*_). pS105 is the control plasmid with a functional **λ** lysis cassette, and is the spanin minus version, *Rz*_*am*_*Rz1*_*am*_. C) MG1655 (*λ900 Rz*_*am*_*Rz1*_*am*_) carrying pRE with ϕKT lysis genes and gp*28*, nonfunctional allele gp*28-HisE17X*, or *RzRz1* were thermally induced and monitored for lysis. Comparison is to uninduced cells. Representative curves from at least two replicates are shown.

To identify these proteins we searched for a segment of the ϕKT genome that would encode all the lysis gene functions. We have previously established a two-plasmid system with the λ lysis cassette (Fig. 1A) cloned under its native late promoter on a medium copy plasmid (pS105) and a second plasmid carrying the gene for the λ late transcription factor Q under inducible control (pQ) (14, 15). Cells carrying these plasmids undergo saltatory lysis if induced during logarithmic phase; experiments with mutants in each of the four lysis genes faithfully replicate the defective lysis phenotypes of the corresponding induced lysogens, including the spherical cell phenotype for null mutants of the spanin subunits (Fig. 2A)(5). Saltatory lysis was also obtained by induction of cells carrying pKT, a plasmid analogous to pS105 with the *27-28-29* gene cluster of ϕKT replacing the λ lysis genes. This indicated that these three genes constitute a complete lysis cassette (Fig. 2B). Moreover, assuming that genes *27* and *29* encode the ϕKT endolysin and holin, respectively, these results also suggest that gp*28* fulfills the OM disruption function provided in λ by the Rz/Rz1 spanin complex. To address this hypothesis we created gene *28* null versions of plasmid pKT and tested them in the same induction system (Fig. 2B). Lysis was abrogated with both nonsense and deletion alleles. In addition, the terminal lysis defective phenotype was spherical cells. Taken together, these results strongly indicate that the holin-endolysin (gp*29* and gp*27*) system of ϕKT achieves the degradation of the PG and that the role of gp*28* is for OM disruption in ϕKT lysis.

To determine if this capacity for OM disruption was specific for the ϕKT holin-endolysin system, we tested gp*28* for its ability to complement the spanin lysis defect in λ inductions (Fig. 2C). Lysogens carrying a λ *Rz*_*am*_*Rz1*_*am*_ prophage and plasmids with either RzRz1 or gene *28* under late promoter control were induced in logarithmic phase and monitored for culture mass and cell morphology. Saltatory lysis was obtained for both constructs, essentially indistinguishable in terms of kinetics and efficiency of the drop in optical density. We conclude that gp*28* does not rely on a special feature of the ΦKT lysis system and, instead, is fully capable of OM disruption even in the λ context.

### The gp*28* disruptin is a cationic membrane-associated polypeptide

Because its primary structure rules out any mechanism requiring a periplasm-spanning complex connecting the IM and OM, as provided by the two-component and unimolecular spanins, we have designated gp*28* as the founding member of a new class of lysis proteins, the “disruptins”. The 56 residue primary structure of gp*28* is marked by a high fraction of charged residues (15 of 56 residues), with a net predicted charge of +7 and three clusters of basic residues at the N-terminus, middle, and C-terminus of the polypeptide (Fig. 3A). Since it is predicted to be a hydrophilic molecule and lacks any predicted membrane localization sequence, yet functions to disrupt the OM, we asked where gp*28* localized during lysis.

**Figure 3.**
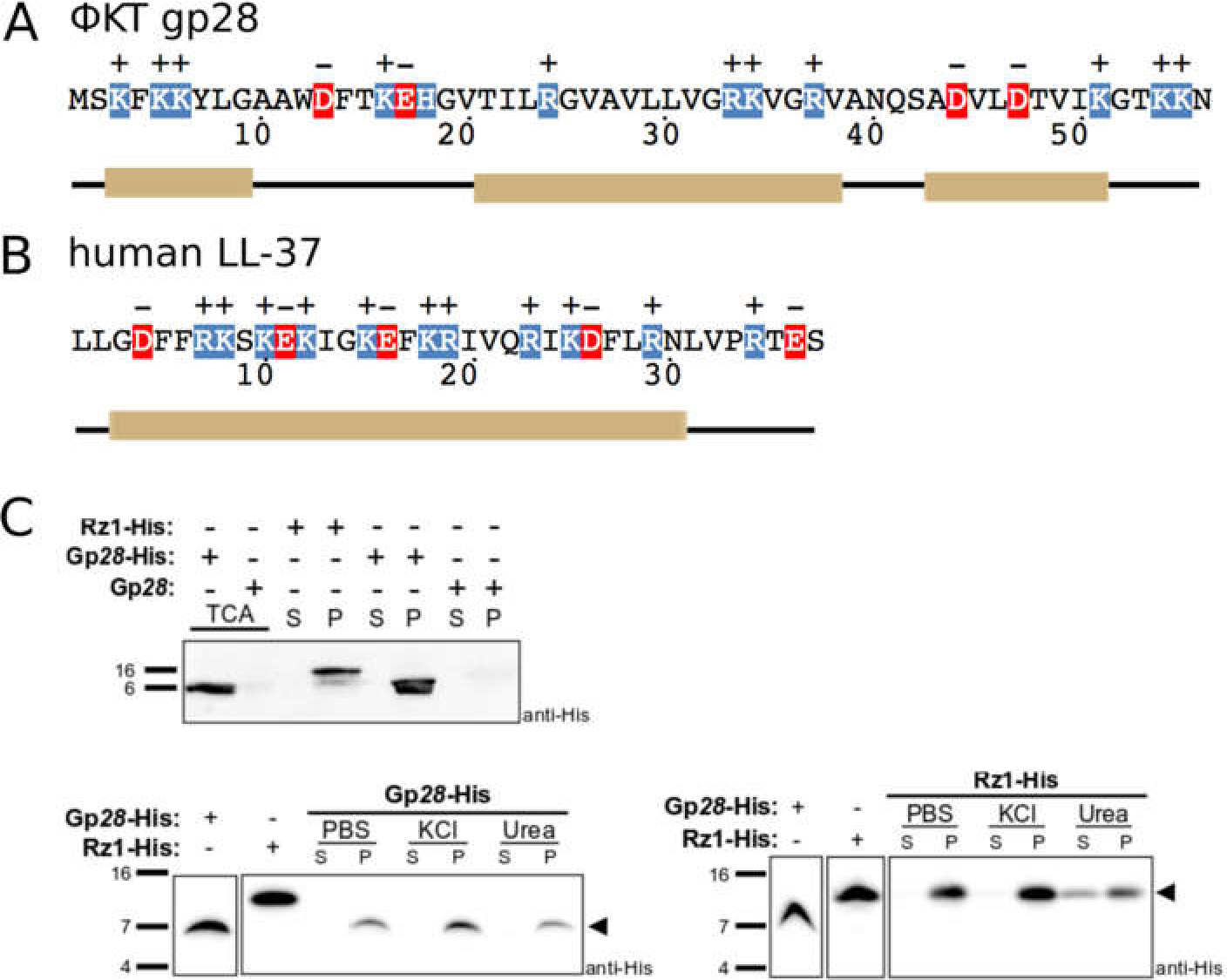
Hydrophobic interactions sequester Gp*28*-His in the membrane. A) The 56 amino acid ϕKT Gp*28* protein sequence with positive (blue) and negative (red) residues highlighted. Alpha-helical content predicted by JPred (33) is indicated below. B) The human LL-37 peptide sequence of 37 residues with positive (blue) and negative (red) amino acids highlighted. Known alpha-helical structure is depicted below. C) Whole MG1655 (λ900 *Rz*_*am*_*Rz1*_*am*_) cells with a pRE plasmid encoding the denoted protein were grown as described previously, collected 55 minutes after induction, and subjected to repeated ultracentrifugation steps to fractionate soluble supernatant (S) and insoluble pellet (P) proteins, followed by TCA precipitation and resolubilization. Immunodetection of the gp*28*-His construct, and its control the Rz1-His construct, is shown by anti-His Western blot. Additionally, the pellet fraction was exposed to PBS, 2M KCl, or 2M urea for solubilization prior to western blotting.

To investigate the subcellular localization of the gp*28* protein, we generated a C-terminal His-tagged version and demonstrated that it retained complementation activity (Fig. 2C). Using this *28-his* allele to complement the lysis defect of λ *Rz*_*am*_*Rz1*_*am*_, we fractionated lysed cells into soluble and membrane fractions. The results show that gp*28*-His is quantitatively associated with the membrane fraction (Fig. 3C). Moreover, only a small fraction of gp*28*-His can be removed from the membrane fraction by urea; indeed gp*28*-His is less susceptible to washing from the membrane-associated state than Rz1, even though Rz1 is a lipoprotein (Fig. 3C). Since gp*28* is synthesized in the cytosol but causes OM disruption, we wanted to know whether it could be detected in both membranes. However, since traditional methods of separating the IM and OM cannot be used on cells that have undergone holin-mediated lysis(6), we used stochastic optical reconstruction microscopy (STORM) to analyze whole cells in which the *28-his* allele was expressed and examined with anti-His antibody (Fig. 4A) (16) (17). Furthermore, to distinguish between inner and outer membrane localization, we used plasmolysis in which the inner membrane retracts from the outer membrane. Thus, inner membrane proteins are contained within “plasmolysis bays” usually located at the cell pole(s) and can be distinguished by a dark spot within the brightfield or phase contrast image of the cell, while outer membranes retain a peripheral localization (18, 19). The results indicate that gp*28*-His is associated primarily with the OM, but up to 20% is located in the IM (Fig. 4B). Because we are imaging the projection of the three-dimensional signal onto a two-dimensional plane, outer membrane signal might be contained within the plasmolysis bays by chance. To account for this, using the same number of molecules present in each cell, we simulated a random distribution of molecules along a cylinder with the same cell width, projected the three-dimensional coordinates onto the two-dimensional imaging plane, and again quantified the fraction of total cellular fluorescence contained in the plasmolysis bays (Fig. 4C). Gp*28*-His distribution is more than two-fold greater than the random simulation control, indicating that greater than half of the signal is true inner membrane localization. We conclude that gp*28*-His is a membrane-bound, cationic polypeptide.

**Figure 4.**
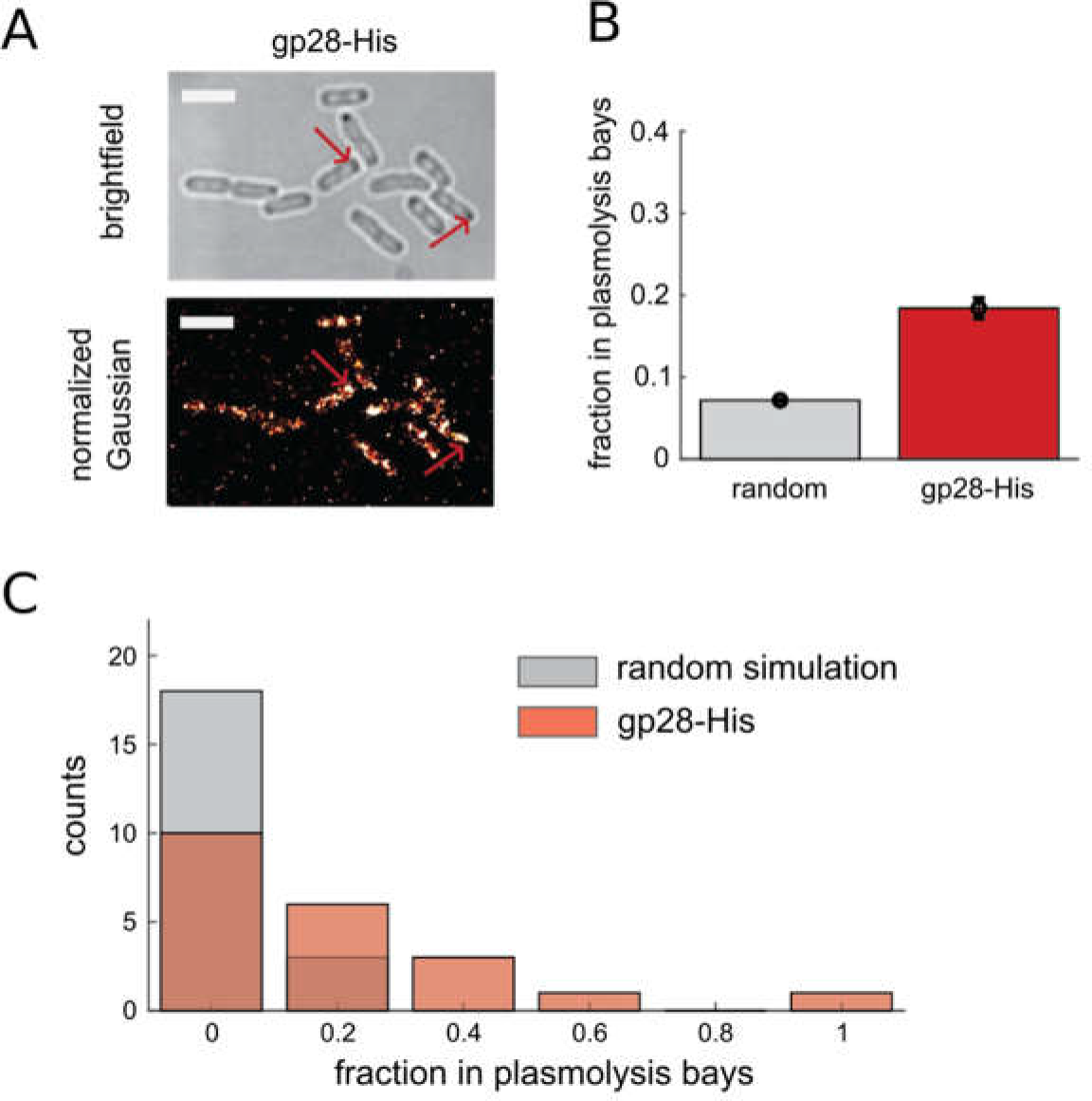
Localization of gp*28*-His in both the inner and outer membranes. MG1655 cells expressing gp*28*-his were fixed and plasmolyzed, followed by fluorescence detection and STORM imaging. A) Example image of plasmolyzed cells; fluorescent images are reconstructions of all detected localizations, red arrows indicate example plasmolysis bays, error bar = 3 microns. B) Quantified fraction of total cellular signal contained within plasmolysis bays, gray bars indicate the signal that contributes by chance from projection of a three-dimensional signal onto a two-dimensional imaging plane; n=21 and error bars are s.e.m. C) Distribution of the fraction of total cellular signal within plasmolysis bays compared to the simulated control.

### Gp*28* can function as the cationic antimicrobial peptide LL-37

The defining characteristics and activity of gp*28*, including its small size, highly cationic primary structure, predominance of predicted alpha helical character, membrane association, ability to cross membranes, and membrane-disrupting function are the hallmarks of the cationic antimicrobial peptides (CAMPs). The human CAMP LL-37 (20, 21) (Fig. 3B), has been the subject of extensive studies in terms of its lethal interaction with bacterial cells, especially in crossing both the OM and IM. The mature polypeptide is 37 aa, has a +6 overall charge and is largely alpha-helical. The CAMPs as a class are thought to disrupt the OM by competing for binding sites for divalent cations in the LPS(22, 23). To assess its capacity to act as a CAMP, we tested a full-length synthetic gp*28* in standard minimal inhibitory concentration (MIC) and minimal bactericidal concentration (MBC) assays. The results show that gp*28* performed at least equivalently to LL-37 against both the native host for ϕKT, 4s, and MG1655, the canonical laboratory *E. coli* strain (Fig. 5).

**Figure 5.**
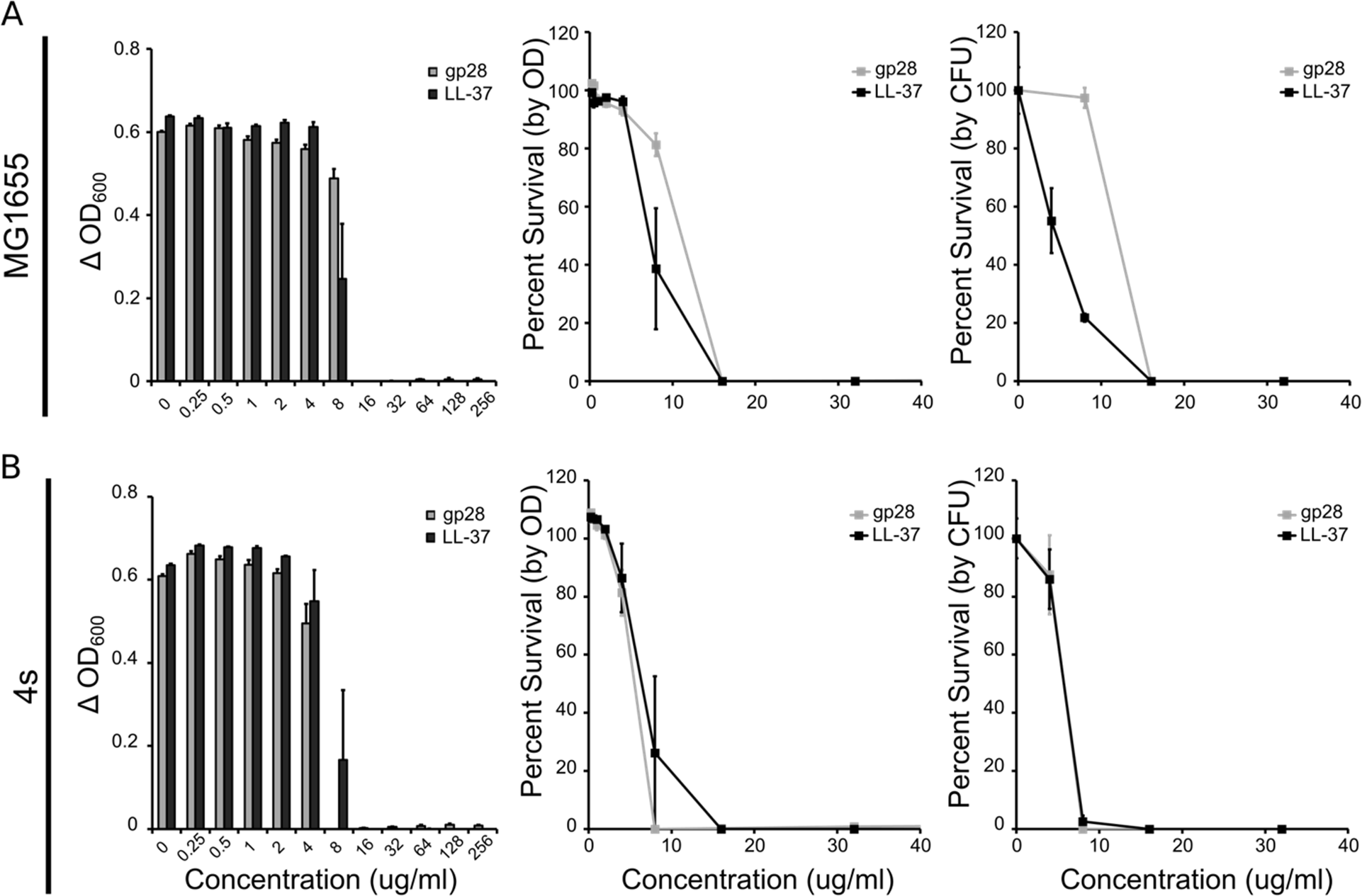
Gp*28* is active as an antimicrobial peptide. *E. coli* A) MG1655 and B) 4s strains were grown in standard MIC and MBC conditions. Cells were treated by exogenous addition of synthesized gp*28* and LL-37 in 10-fold decreasing amounts for 16 hours. The change in optical density, and percent survival as a function of both optical density and viable cells is plotted as a function of peptide concentration.

## Discussion

### The first disruptin

OM disruption function is required for lysis in phage λ and is provided by its Rz/Rz1 two-component spanin system. The simplest notion is that OM disruption is required for all phages of Gram-negative hosts. Here, we report that for a coliphage where we could rule out the presence of any kind of lipoprotein gene (and thus unimolecular or two-component spanins), a new type of OM-disruption protein, the disruptin, was found. This protein, ϕKT gp*28*, fully complements the lysis defect associated with the absence of spanin function in the canonical holin-endolysin system. Remarkably, gp*28* has many features that are hallmarks of CAMPs like the well-studied human cathelicidin LL-37, including small size, a sequence rich in basic residues and dominated by predicted alpha-helical structure, and the ability to bind to and pass through membranes. Moreover, synthetic gp*28* exhibited antimicrobial activity comparable to LL-37 in standard MIC and MBC assays.

Gp*28* has no homologs in the database, except for a nearly identical protein in a very closely related phage PGT2 (NCBI Accession: <u>ATS92463.1</u>). We expect other spaninless phages that infect Gram-negative hosts also to encode disruptins with characteristics that match the CAMP or other AMP families. The task of identifying disruptins bioinformatically will be challenging, especially in uncharacterized or larger phage genomes with more than one predicted small, cationic, alpha-helical protein that may or may not be near other lysis genes. In a preliminary analysis of the 91 spaninless genomes, 528 candidates meet an arbitrary cutoff of <80 aa and a minimum +4 net charge. Although choosing candidates in this way is noisy, the availability of the facile and rigorous RzRz1-complementation system used here makes this approach workable. Work along these lines is underway and could elucidate how general the disruptin solution is for OM destruction during phage lysis.

### How does the disruptin function for OM disruption?

Gp*28* was detected only in the particulate fractions, indicating that the gp*28* peptide has a strong affinity to membranes. STORM analysis further revealed that the disruptin is also found in the OM, even when expressed in the absence of other lysis proteins. The simplest notion is that after synthesis in the cytosol, gp*28* binds to the IM, after which it passes through to the OM where it binds preferentially. Both the preferential localization of gp*28* to the OM and its OM disruption function may be explained by the ability of CAMPs to compete for the divalent-cation binding sites in the LPS layer (17, 22). Removal of a fraction of the stabilizing Mg^++^/Ca^++^ ions resident in the phosphate-rich core LPS would likely reduce the tensile strength of the OM. Even a small reduction in the OM strength would limit the load-bearing capacity required for resisting the turgor pressure at the last step of the lysis pathway. In other words, after endolysin-mediated destruction of the PG, a weakened OM would be susceptible to rupture.

### Disruptin function in the context of holin control

Holins control the timing of lysis and thus the length and fecundity of the infection cycle. The timing is thought to reflect a characteristic mass-action property of holins, which accumulate harmlessly in the IM until reaching an allele-specific critical concentration and triggering to form lethal holes. This preserves the integrity of the IM and thus maximal biosynthetic capacity throughout the viral assembly phase of the infection cycle. Moreover, since single missense changes throughout the length of the holin can cause dramatic and unpredictable alteration of the critical concentration, a phage can rapidly evolve to drastically different lengths of the infection cycle. Theoretical and experimental analyses have suggested that environmental factors can favor shorter or longer infections, especially the average density of available hosts. Both two-component and unimolecular spanin proteins accumulate in the envelope throughout the morphogenesis phase and thus are capable of subverting holin-mediated control of the lysis pathway. However, both types of spanins are blocked from lethal function by being encaged in the meshwork of the PG network, which in turn is maintained until holin-controlled activation of the endolysin. From this perspective, the use of the disruptin for the last step in the lysis pathway is surprising, in that it not only pre-localizes to the IM and the OM irrespective of holin or endolysin function but also offers no obvious feature that would confer sensitivity to holin control. Moreover gp*28* appears to have bactericidal capacity essentially indistinguishable from that of the well-studied CAMP LL-37. CAMPs are known to permeabilize both the IM and OM and cause collapse of the PMF. This seems even more problematical for the control of lysis timing, since holins can be prematurely triggered by even partial reduction of the PMF. The key to compatibility of the disruptin with holin control likely resides in quantitative considerations. Membrane permeation typically requires binding of >10^6^ CAMP molecules to the bacterial cell (24). However, the levels of expression of the lysis proteins in λ infections is on the order of 10^3^ to 10^4^ molecules at the time of lysis. Although the level of expression of gp*28* in ϕKT infections has not yet been directly measured, it supports lysis when expressed from the λ late promoter under its native and unexceptional Shine-Dalgarno ribosome binding site. We suggest that at the levels produced during the infection cycle the disruptin neither affects the PMF as it binds and passes through the IM, nor significantly permeabilizes the OM as it binds the LPS and displaces a small fraction of the phosphate-neutralizing divalent cations. Accordingly the effects of disruptin binding are not realized until the entire load of the cell turgor pressure is transferred to the OM by the endolysin-mediated degradation of the PG. Experiments using the non-invasive flagella-tethered cells to quantify the PMF during gp*28* expression can be used to test this model(25, 26).

This raises an interesting question for how disruptins may be compatible with holins that are programmed to trigger lysis at later times after induction. Genetic analysis has shown that single base changes in the holin gene alter the time at which lysis is initiated. This results in the shortening or extension of the morphogenesis period and significantly affects the amount of phages that are released at the time of lysis. Importantly, this has been linked to how phages can respond to changes in the environment of the host. That is, a shorter lifecycle provides fitness in a host-rich environment whereas a longer lifecycle is favored in a host-depleted setting (27, 28). We would expect that the continued accumulation of the disruptin would abrogate the PMF, which would trigger the holin to initiate lysis prematurely. This could be tested with the disruptin under control of a tunable expression system in the presence of holin-endolysin systems programmed to initiate lysis at different intervals. If this were found to be true, a consequence would be to the fecundity of disruptin lysis systems, which would then be limited to initiating lysis before the accumulating disruptin affects holin control of lysis timing. This may explain in part why spanins are more abundant even though they require more genetic space

## Materials and Methods

### Strains, phages, plasmids, and primers

The strains, phages, and plasmids used in this study are presented in **Table 1**. The primers used for this study are summarized in **Table 2**.

**Table 1.**
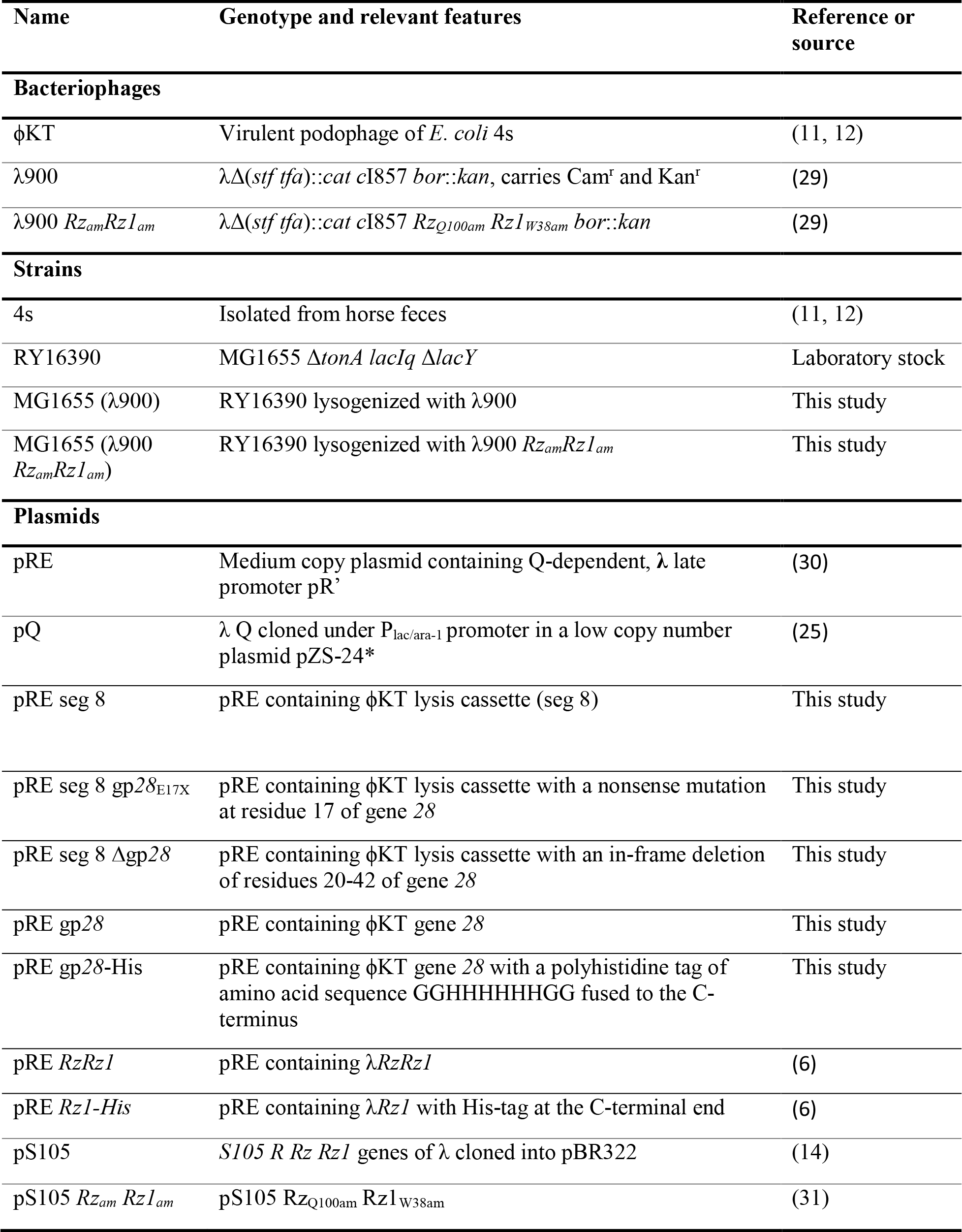
Strains, phages, and plasmids.

**Table 2.**
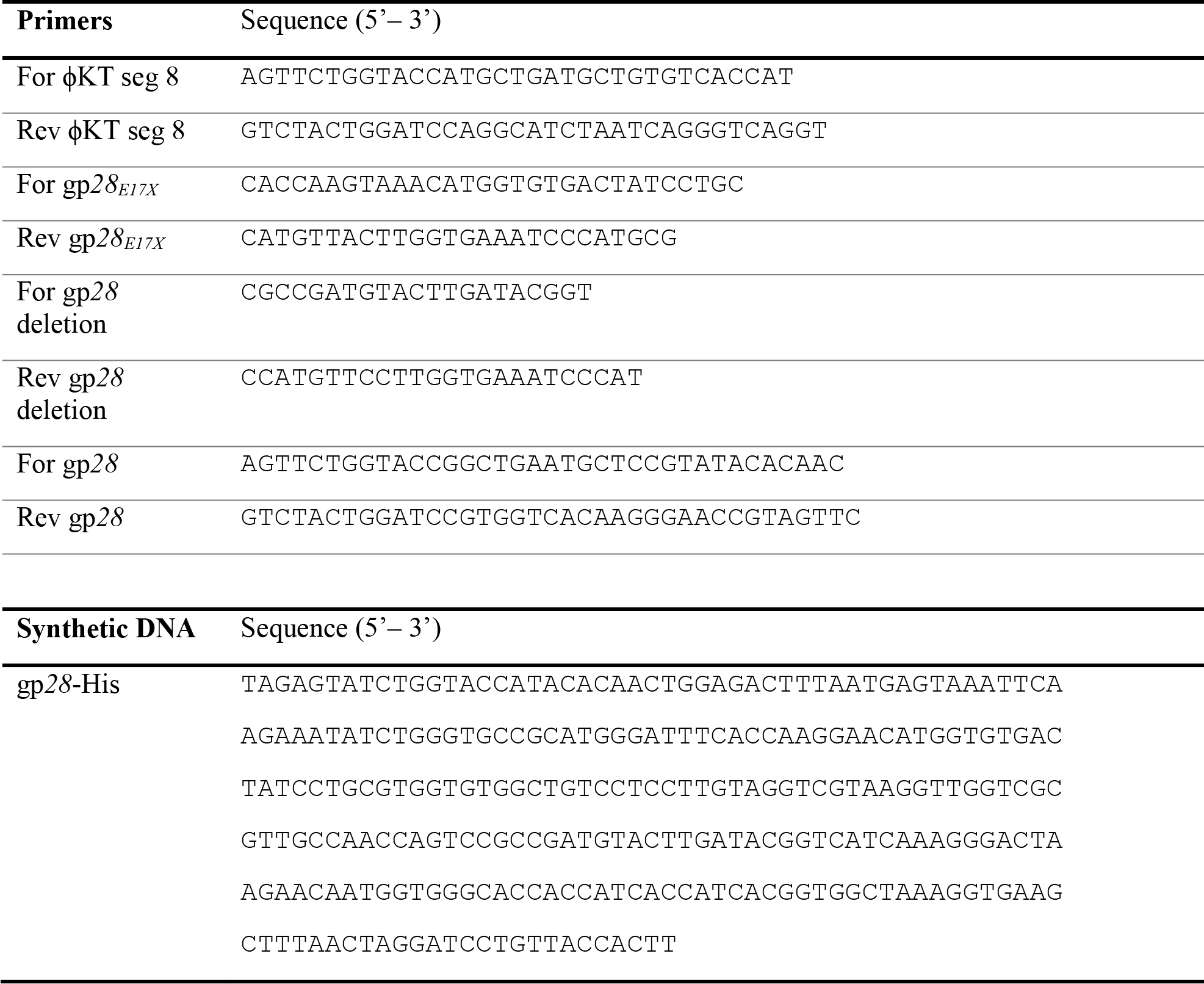
Primers and DNA constructs.

### DNA manipulation

The pRE seg 8 plasmid was constructed by PCR amplifying the ΦKT lysis cassette with the For ΦKT seg 8 and Rev ΦKT seg 8 primers. These primers contain KpnI and BamHI overhangs and were designed to amplify the section of the ΦKT genome from nucleotide 36,581 to 37,741 (GenBank Accession: NC_ 019520.1). The gel-purified PCR product was double digested with the corresponding restriction enzymes and ligated into pRE using T4 DNA ligase. The pRE gp*28* plasmid was constructed by a similar method, the only modification being that primers For gp*28* and Rev gp*28* were used to amplify from nucleotide 37,067 to 37,323 in the ΦKT genome. Quick change modifications were performed as described previously (8).

### Bacterial growth, induction, and lysis monitoring

Bacterial cultures were grown, induced, and monitored for lysis as described previously (8).

### ΦKT infection

Bacterial cultures were grown at 37°C to an A_550_ of ~0.3. Then the cultures were put on ice, and phage were added to the indicated samples at a multiplicity of infection (MOI) of 30. After 90 minutes on ice, the cultures were again incubated at 37°C and monitored for lysis.

### Phase contrast microscopy

A 1.5 μL culture sample was applied to a glass slide and covered with a coverslip. The sample was imaged immediately using a plan-apochromat 20x/0.8 Ph2 objective installed on a Zeiss Axio Observer 7 inverted microscope. All image processing was done using the Zen 2.3 software.

### Subcellular localization

Cells were induced at 0.25 A_550_ and 10 mL samples were collected 55 minutes after induction, after the culture had cleared. Cells were placed on ice, and 1 μL of protease inhibitor cocktail (Sigma-Aldrich, P8465) was added to each 10 mL sample. This lysate was spun down at 6,000 x *g* for 5 minutes at 4°C to clear unlysed cells. Then, three 3 mL aliquots of supernatant were taken from each 10 mL sample and spun at 100,000 x *g* for 60 minutes at 4°C in a TLA 100.3 rotor using an Optima MAX-XP ultracentrifuge. After the centrifugation, the supernatant was removed, and 500 μL of 1x PBS, pH 7.4 was added to the pellet. The pellet was incubated in the PBS at 4°C overnight for resuspension. An additional 2.5 mL of PBS was added to each sample before a second centrifugation was performed at 100,000 x *g* as described above. Next, 500 μL of 2 M KCl, 1x PBS, or 2 M urea was added and samples were resuspended by incubating cold overnight. Finally, 2.5 additional mL of the same solution was added to each sample and a final ultracentrifugation was carried out as described above. After centrifugation, 1 mL of the supernatant was collected and a proteins were precipitated with 10% TCA as previously described (4, 32). Both the ultracentrifugation and TCA precipitate pellets were resuspended in 1x sample loading buffer at 100 μL per 1 OD unit as measured at 40 minutes after induction of the original bulk culture. After boiling for 5 minutes, samples were electrophoresed and detected by blotting as described below.

### SDS-PAGE and western blotting

Samples were collected and prepared for Western blotting as described previously (8). After preparation, the samples were resolved on a Novex 10-20% Tricine SDS-PAGE gel (Thermo Fisher, Waltham, MA). Gel transfer and immunodetection were done using the iBlot and iBind systems (Thermo Fisher, Waltham, MA) using the manufacturer’s recommended protocol. For the primary antibody step, mouse anti-His antibody (Thermo Fisher, Waltham, MA) was diluted 1:2000 in 1x iBind solution. A goat anti-mouse-HRP (Thermo Fisher, Waltham, MA) secondary antibody diluted 1:1000 in 1x iBind solution was used for detection. Chemiluminescence scanning was performed on an Amersham Imager 600 RGB (GE, Pittsburgh, PA).

### Localization of gp*28*-His by STORM imaging

Wild-type MG1655 cells carrying pQ and pRE gp*28-His* were induced with 1 mM IPTG after reaching an OD550 of 0.2. After one hour growth, cells were harvested by centrifugation and plasmolysed by resuspension in plasmolysis buffer (15% sucrose, 25mM HEPES, 20mM sodium azide). Cells were then fixed for 15 min at room temperature in fixation buffer (1x PBS, 2.6% paraformaldehyde, 0.8% glutaraldehyde, washed twice in 1x PBS, then rotated at 37°C for 30 minutes in PBS + 10% goat serum (Sigma Aldrich, Saint Louis, MO). A 1:150 dilution of anti-His antibody labelled with AF647 (Thermo Fisher, Waltham, MA) was then added to cells, and cells were rotated at 4°C in the antibody mixture overnight (~16 hours). Cells were then washed four times in 1x PBS + 0.1% Tween-20 solution, with a final resuspension in 1x GTE buffer. To visualize the localization of labelled gp*28*-His in cells, the antibody-labelled cells were adhered to coverslips treated with 0.1% poly-L-lysine (Ted Pella, Redding, CA) and imaged on an Olympus IX81 inverted microscope, with an 100x oil-immersion objective with NA of 1.45 using a highly inclined thin illumination imaging scheme. Fluorescent molecules were imaged using a high power (100 mW) of 647 nm laser light (Coherent), with images acquired every 10 ms, with a total of 15,000 images collected per region imaged. Emitted fluorescence was first filtered through an emission filter (705/50, ThorLabs) and then collected on an Andor emCCD camera with 300x gain.

For each region imaged, individual cells were manually identified and cropped out using the ImageJ ROI Manager; both brightfield and fluorescence tiff stacks were cropped for each cell and saved as individual substacks. These individual substacks were then imported into MATLAB and using custom-built software, plasmolysis bay and cell ROIs were manually identified using the brightfield images of the cells. Using these ROIs, the relative fluorescence fraction of signal in plasmolysis bays versus the entire cell was calculated. Because the images were acquired in two-dimensions, some signal from the outer membrane could be acquired in the ROI of the plasmolysis bay by chance. To account for this, for each cell, we simulated what the distribution of molecules would be for randomly deposited molecules onto a three-dimensional cylinder the same width as the cell, using the number of molecules that were present for that particular cell. The fraction of signal in the plasmolysis bay ROI vs the entire cell ROI was again calculated for this random simulation. The random simulation signal fraction represents the average fraction that is contributed by outer membrane signal to the signal in the plasmolysis bays. We compared the mean random simulation fraction to the mean acquired signal for gp*28*-His, as well as the distribution of fractions for the random simulation versus the cellular signal.

### Minimal inhibitory concentration (MIC) and minimal bactericidal concentration (MBC_99_) of peptides gp*28* and LL-37

Peptides gp*28* and LL-37 were dissolved in ddH2O to give a stock concentration of 5.12 mg/ml and aliquots were stored at −80°C. The maximum concentration of peptides used for the assays was 256 µg/ml. For each assay, primary cultures of MG1655 or 4s were started from a −80°C stock and grown overnight in Mueller Hinton (MH) media at 37 °C shaking. A secondary culture started in late log phase, with an optical density (OD_600_) of 1.5 to 2.0 was used to prepare the starting inoculum in MH media with 0.001% acetic acid and 0.02% BSA to give a calculated OD of 0.001, which corresponded to about 5 × 10^5^ colony forming units (CFU/ml) of bacteria. Using 96-well polypropylene, non-pyrogenic cell culture plates (Corning Incorporated, Corning, NY), 100 µl of 5 × 10^5^ CFU/ml bacteria were added in triplicate for each peptide. Each antimicrobial agent was assayed at 11 different concentrations, in two-fold serial dilutions. They were grown in a humidity controlled incubator at 37 °C for 16 hours. OD_600_ was measured using a PerkinElmer Envision 2104 multi-label plated reader (Perkin Elmer, Waltham, MA) at the start of the assay and at 16 hours. After 16 hours, bacteria were plated for colony forming unit (CFU/ml) counts in MH agar plates. Percent survival was determined for each peptide concentration using the OD_600_ and CFU/ml. OD_600_ and CFU/ml were compared to the bacteria only control at 16 hours. The MIC and MBC_99_ values were calculated relative to the starting CFU/ml. MIC was defined as the minimal concentration of the peptide needed to inhibit growth of the starting culture of bacteria (≤100%). MBC_99_ was defined as the minimal concentration of the peptide needed to kill 99% of the starting culture of bacteria (≤1%). These assays were done in triplicate.

## Acknowledgments

This work was supported by Public Health Service grant GM27099, National Science Foundation Award number HRD-1304975, two Beckman Scholars Program Awards from the Arnold and Mabel Beckman Foundation, and by the Center for Phage Technology at Texas A&M University, jointly sponsored by Texas AgriLife. We thank Young lab members, past and present, for their valuable input during the course of this study.

